# Developmental exposure to the Parkinson’s disease-associated organochlorine pesticide dieldrin alters dopamine neurotransmission in α- synuclein pre-formed fibril (PFF)-injected mice

**DOI:** 10.1101/2023.06.21.545967

**Authors:** Sierra L. Boyd, Nathan C. Kuhn, Joseph R. Patterson, Anna C. Stoll, Sydney A. Zimmerman, Mason R. Kolanowski, Joseph J. Neubecker, Kelvin C. Luk, Eric S. Ramsson, Caryl E. Sortwell, Alison I. Bernstein

## Abstract

Parkinson’s disease (PD) is the most common movement disorder and one of the fastest-growing neurological diseases worldwide. This increase outpaces the rate of aging and is most rapid in recently industrialized areas, suggesting the role of environmental factors. Consistent with this, epidemiological studies show an association between exposure to persistent organic pollutants and an increased risk of PD. When combined with post-mortem analysis and mechanistic studies, a role for specific compounds, including the organochlorine pesticide dieldrin, emerges. In mouse models, developmental dieldrin exposure causes male-specific exacerbation of neuronal susceptibility to MPTP and synucleinopathy. Specifically, our novel two-hit model combining developmental dieldrin exposure with the α-synuclein (α-syn) pre-formed fibril (PFF) model showed a male-specific exacerbation of PFF-induced increases in striatal dopamine (DA) turnover and motor deficits on the challenging beam 6 months post-PFF injection in male offspring developmentally exposed to dieldrin. Here, we hypothesized that alterations in DA handling contribute to the observed changes and assessed vesicular monoamine transporter 2 (VMAT2) function and DA release in this dieldrin/PFF two-hit model. Female C57BL/6 mice were exposed to 0.3 mg/kg dieldrin or vehicle every 3 days, starting at 8 weeks of age by feeding and continuing throughout breeding, gestation, and lactation. Male offspring from independent litters underwent unilateral, intrastriatal injections of α-syn PFFs via stereotaxic surgery at 12 weeks of age and DA handling was assessed 4 months post-PFF injection via vesicular ^3^H-DA uptake assay and fast-scan cyclic voltammetry (FSCV). We observed no dieldrin-associated change in VMAT2 activity, but a dieldrin-induced increase in DA release in striatal slices in PFF-injected animals. These results suggest that developmental dieldrin exposure alters the dopaminergic response to synucleinopathy-triggered toxicity and supports our hypothesis that alterations in DA handling may underly the observed exacerbation of PFF-induced deficits in motor behavior and DA turnover.

**Graphical Abstract:** 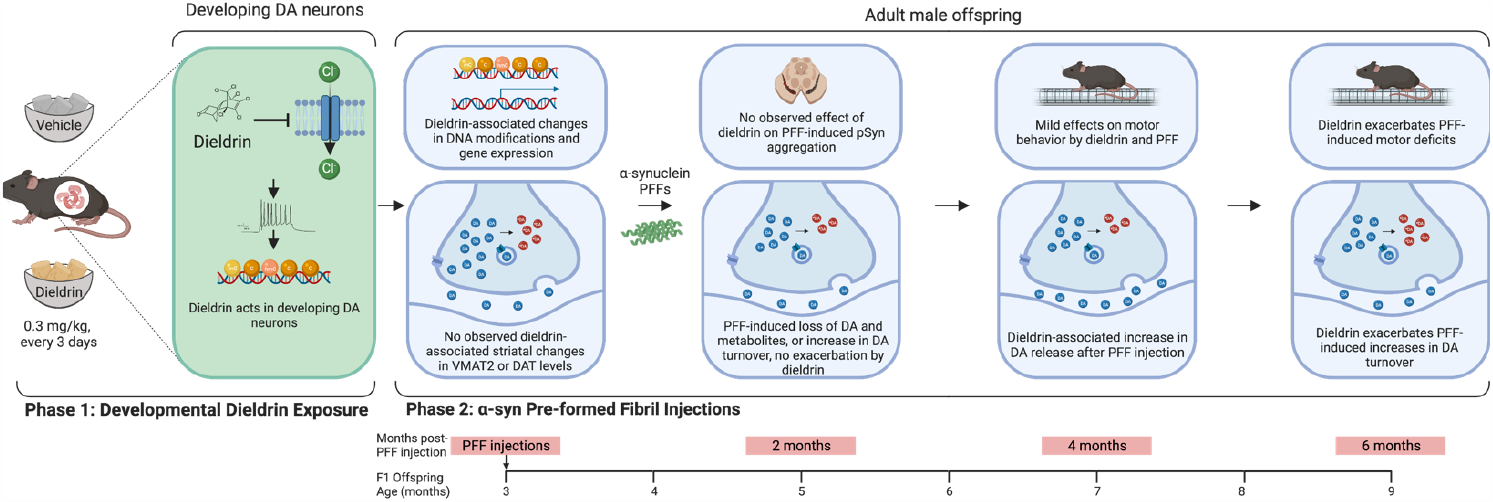

## Introduction

Parkinson’s disease (PD) is a multi-system disorder pathologically defined by the degeneration of dopaminergic neurons in the nigrostriatal pathway and the formation of α-synuclein (α-syn)-containing Lewy bodies. PD is the most common movement disorder, the second most common neurogenerative disease, and one of the fastest growing neurological diseases.^1^ From 1990 to 2016, the prevalence of PD has more than doubled globally.^2^ In addition, a recent study suggests that PD incidence in the US is 50% higher than previously estimated, with 90,000 diagnoses per year^3^. Of relevance to this work, the authors reported PD incidence rates higher in certain geographic areas including the “Rust Belt,” a region with a history of heavy industrial manufacturing. This is consistent with epidemiological research that shows an association between increased risk of PD and environmental factors associated with industrialization, including heavy metals, solvents, and pesticide exposures.^1,2,4–17^ When combined with post-mortem analysis and mechanistic studies, a role for specific compounds in PD emerges. One such compound is dieldrin, an organochlorine pesticide that is associated with an increased risk of PD in both epidemiological and mechanistic studies. ^12,18–21^

We previously developed a two-hit model that combines a developmental dieldrin exposure paradigm with the α-synuclein (α-syn) pre-formed fibril (PFF) synucleinopathy model to explore the mechanisms that underly exposure-related increases in PD susceptibility.^22–24^ In this model, dams are fed dieldrin throughout mating, gestation, and lactation and F1 pups are assessed for toxicity in PD models at 12 weeks of age. ^24,25^ Research in our lab and others has shown that this developmental dieldrin exposure paradigm induces stable alterations in the DA system that increase susceptibility to subsequent exposure to both PFF-induced synucleinopathy and MPTP in male, but not female, offspring.^24–26^ The exacerbation of synucleinopathy-induced behavioral effects is specific to male mice, with female mice showing less nigral TH loss and no behavioral deficits ^24,25^.

This sex specificity is consistent with the reduced incidence of PD and severity of disease course in human females, known sex difference in dopaminergic vulnerability to parkinsonian toxicants, and our previously reported sex-specific epigenetic effects of developmental dieldrin exposure.^27–38^ Specifically, in male F1 offspring, we found dieldrin-induced exacerbation of synucleinopathy-induced deficits in striatal DA turnover and motor deficits on challenging beam 6 months after PFF injection.^24^ However, exposure does not affect PFF-induced aggregation of α-syn at 1 or 2 months post-PFF injection, overall loss of striatal DA or the DA metabolites, DOPAC and HVA, at 2 or 6 months post-PFF injection, or loss of nigral tyrosine hydroxylase (TH)- or NeuN-positive neurons at 6 months post-PFF injection.

Proper packaging of DA into synaptic vesicles and trafficking of vesicles to the active zone are critical for DA neurotransmission.^39^ In addition, while DA is stable within the acidic environment of a synaptic vesicle, cytosolic DA is metabolized to DOPAC and HVA or broken down by auto-oxidation producing toxic products. As a result, excessive levels of cytosolic DA are neurotoxic to DA neurons and mishandling of DA can directly affect α-syn aggregation and induce PD-like pathology.^39–51^ Insults that disrupt DA packaging and vesicular function impair DA release and increase toxicity via elevated cytosolic DA and the relative expression or activity of DAT (dopamine transporter) to VMAT2 (vesicular monoamine transporter 2) has been suggested as a determinant of neuronal vulnerability in PD.^39,50,51^ Based on the observed exacerbation in PFF-induced increases in striatal DA turnover by dieldrin, we hypothesized that dieldrin-induced alterations in DA packaging and synaptic vesicle dysfunction contribute to the observed changes in PFF-induced animals.

To test this, we assessed VMAT2 function by vesicular uptake assay and DA release by fast-scan cyclic voltammetry (FSCV) in the dieldrin/PFF two-hit model 4 months post-PFF injection in male F1 offspring developmentally exposed to dieldrin. Testing these at 4 months allowed us to capture changes in the striatal synapse prior to significant nigrostriatal degeneration; assessing these at 6 months when degeneration of striatal terminals and nigral cell bodies is already pronounced would test mainly the effects of degeneration, rather than the dysfunction that precedes it. While we did not detect a change in VMAT2 uptake velocity in PFF-injected mice via vesicular ^3^H-dopamine uptake assays as predicted, we observed a dieldrin-related increase in DA release using FSCV. Therefore, our data show that developmental dieldrin exposure modulates PFF-induced responses in DA packaging and synaptic vesicle function. These observed increases in DA release and uptake velocity in PFF-injected animals are consistent with our previously observed increase in DA turnover in PFF-injected animals developmentally exposure to dieldrin.^24^

## Materials and Methods

### Animals

Male (11 weeks old) and female (7 weeks old) C57BL/6 mice were purchased from Jackson Laboratory (Bar Harbor, Maine). Animal husbandry and colony maintenance was completed as previously described. ^24,38^ All procedures were conducted in accordance with the National Institutes of Health Guide for Care and Use of Laboratory Animals and approved by the Institutional Animal Care and Use Committee at Michigan State University.

### Dieldrin exposure paradigm

Dosing was carried out as previously described.^24,38^ Adult C57BL/6 (8-week-old) female animals were treated throughout breeding, gestation, and lactation (Figure 1A). Mice were administered 0.3 mg/kg dieldrin (ChemService) dissolved in corn oil vehicle and mixed with peanut butter pellets every 3 days. ^24,38,52^ Control mice received an equivalent amount of corn oil vehicle in peanut butter. This dose was based on previous results showing low toxicity, but clear developmental effects. ^25^ Four weeks into female exposure, unexposed C57BL/6 males (8–12 weeks old) were introduced for breeding. Offspring were weaned at 3 weeks of age and separated by litter and by sex, with 2-4 animals per cage (Figure 1B). At 12 weeks of age, male offspring from independent litters were selected for PFF or monomer injection. This time point was chosen based on previous results demonstrating increased neuronal susceptibility to this age. ^24,25^This developmental dieldrin dosing paradigm has been previously used in our lab to study the role of epigenetics and the effects on the development of synucleinopathy. ^24,38^

**Figure 1.**
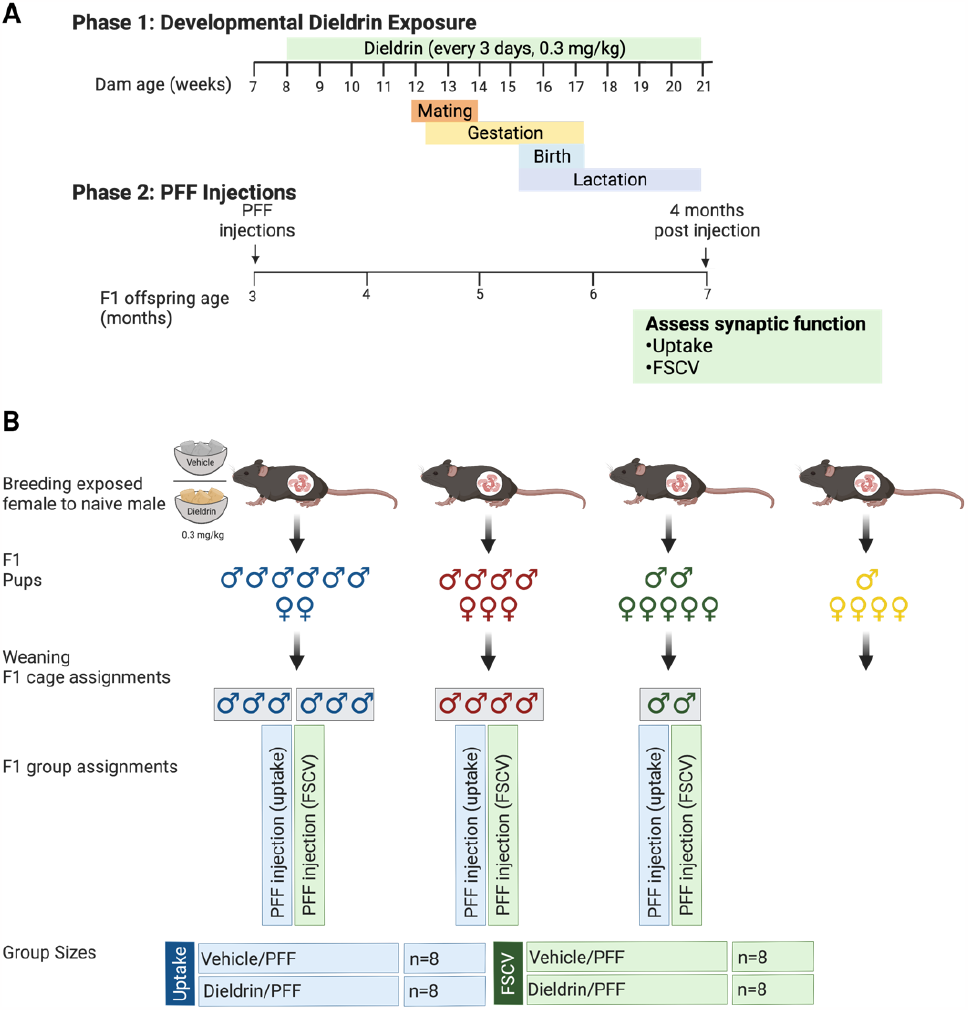
Experimental design including dosing schedule, weaning strategy, cage and group assignments. A) Timeline of dieldrin-PFF two-hit model: At 8 weeks of age, female C57BL/6 mice dieldrin exposure began via oral administration of 0.3mg/kg dissolved in corn oil and injected into peanut butter pellets. At 12 weeks of age, mating began, and exposure continued through weaning of pups. F1 pups were weaned 3 weeks after birth and separated by litter and sex (2-4 animals per cage). At 3 months of age, male pups underwent intrastriatal injections of PFFs and were individually housed after surgery. B) Cage, group assignments, and group numbers: Male F1 offspring (F1) that underwent intrastriatal PFF-injections were assigned to endpoints such that every animal for each endpoint came from an independent litter. The Fourth F1 litter is an example of a litter excluded from endpoint assignments due to individual housing.

### Preparation of α-synuclein PFFs and fibril size verification

Recombinant mouse α-syn monomers and PFFs were provided by the Luk lab, stored at -80 °C, and prepared as previously described. ^22,53^ Over 500 fibrils were measured to determine the average fibril length of 45.06nm +/- 14.7nm (Figure 2).Fibril length was assessed before and after surgeries to ensure that fibrils did not re-aggregate over the duration of the surgies. All measurements were performed with ImageJ. ^54^

**Figure 2.**
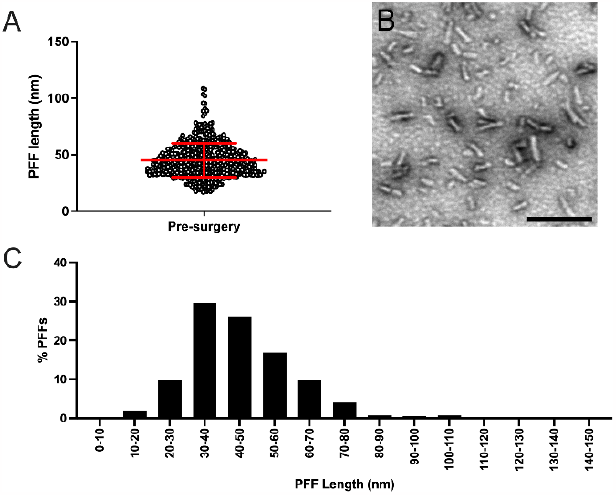
Verification of α-syn PFF size. A) PFF length distribution determined via TEM. Each point represents a measured fibril length, the error bars denote standard deviation. B) Representative TEM image of sonicated fibrils. C) Frequency distribution of PFF lengths post-sonication.

### Intrastriatal injection of α-syn PFFs

At 12 weeks of age, animals received unilateral intrastriatal PFF injections according to their cage and group assignment (Figure 1B). Surgeries were performed as previously described. ^23,24^ Mice received a total of 5 µg of PFFs (2.5 µL injection of 2 µg/µL PFFs) and received a single intrastriatal injection (anterior-posterior (AP) +0.2, medial-lateral (ML) +2.0, dorsal-ventral (DV) -2.6) with a flow rate of 0.5 μl/ml. Post-surgery, mice received 1mg/kg of sustained release buprenorphine by subcutaneous injection and were monitored closely until they recovered from anesthesia. In the three days following recovery, animals were monitored daily for adverse outcomes. A small subset of animals (n=2 for FSCV and an n=4 for uptake assays) received α-syn monomer injections to as a negative control to ensure that there were no effects of surgery itself. Animals were singly housed following surgeries for the duration of the experiment, consistent with our previous study. ^24^

### Vesicular ^3^H-dopamine uptake

Animals were euthanized by cervical dislocation and hemisected. Half of the brain from each group was homogenized for each statistical n, and vesicular DA uptake was performed as previously described. ^44,48,55,56^ Data was normalized to protein level determined by BCA assay and expressed as pmol DA/mg protein/minute.

### Fast-scan cyclic voltammetry

Animals were euthanized by cervical dislocation and brains were sectioned in oxygenated, 4°C artificial cerebrospinal fluid (aCSF) at 300 µm thick using a vibratome (Campden Instruments 5100mz-Plus)^57^. FSCV was carried out in the lateral, dorsal striatal sections as previously described. ^48,57–61^

Carbon fiber glass microelectrodes were constructed using a vacuum to pull carbon fiber through a glass capillary tube, pulled using a horizontal electrode puller, broken, and sealed with paraffin^59^. Microelectrodes were cycled for at least 15 minutes prior to recording at a frequency of 60Hz, then cycled at 10Hz until stable. ^59,60,62,63^ Carbon fiber microelectrodes were calibrated using a pipette-based calibration system by adding a dilute stock DA solution to a buffer and measuring the oxidation and/or reduction ^60^. All cycling and recordings occurred with a triangle waveform (−0.4 to 1.3V to -0.4V; 400V/s 10Hz). ^48,58,59,64^ Dopamine release was elicited with a bipolar twisted electrode (PlasticsOne) and a 350µA, 4ms monophasic optically isolated stimulus pulse (Neurolog NL800). Data was collected and analyzed using Demon Voltammetry and Analysis Software (Wake Forest Innovations). ^61^ A five-recording survey of two different dorsal striatal release sites per hemisphere in 2 different slices was taken for each animal with a 5-min rest interval between each stimulation. ^48,58^ Peak Dopamine, upward velocity (DA release), downward velocity (uptake; a Vmax estimate), and tau (uptake; a Km estimate) were calculated for each recording (Figure 5A). ^58^ Ipsilateral values were normalized to contralateral values to account for animal-to-animal variability.

### Immunohistochemistry

The rostral remainder of the brains used for FSCV were immersion fixed in 4% paraformaldehyde for 24 hours and placed into 30% sucrose in PBS at 4°C for immunohistochemistry. Fixed brains were frozen on a sliding microtome and sliced at 40 μm coronally. Free-floating sections were stored in cryoprotectant (30% sucrose, 30% ethylene glycol, 0.05 M PBS) at -20 °C. A 1:6 series of the entire rostral portion of the brain was used for staining and two nigral sections per animal were selected for imaging and quantification. Nonspecific staining was blocked with 10% normal goat serum, and sections were then incubated overnight in appropriate primary antibodies in TBS with 1% NGS/0.25% Triton X-100 followed by appropriate secondary antibodies for 2 hours (Table 1). Slides were cover-slipped with VECTASHIELD Vibrance Antifade Mounting Medium (VectaLabs) with DAPI and imaged on a Zeiss AxioScan 7 Digital Slide Scanning Microscope. Analysis of pSyn counts for immunohistochemistry was completed using the object colocalization module in the HALO Image Analysis Platform (Indica Labs). The SNpc was manually traced as the region of interest based on TH staining and pSyn positive objects within this region were identified on two sections per animal from the same level.

**Table 1:**
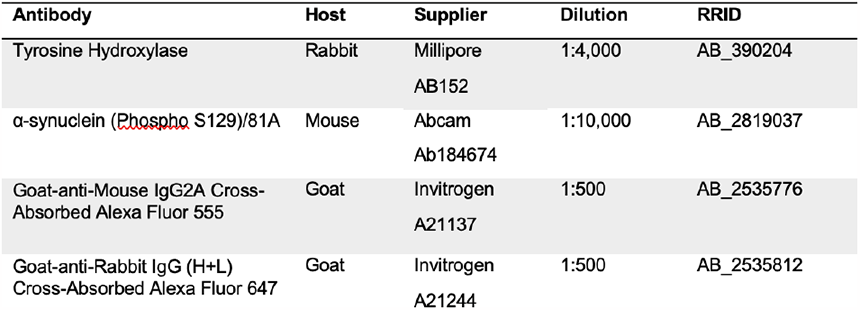
Antibodies used for Immunohistochemistry.

### Data Analysis and Statistics

Statistical analysis and graphing were performed using GraphPad Prism 9. Vehicle-exposed animals injected with PFFs (Vehicle/PFF) and dieldrin-exposed animals injected with PFFs (Dieldrin/PFF) were compared with two-tailed, unpaired-t-tests. All data are shown as mean +/- SD and cutoff for statistical significance was p<0.05. Monomer injected animals (Vehicle/Monomer, Dieldrin/Monomer) were used as controls to ensure there were not effects of surgery on its own, but these were not included in the statistical analysis, consistent with our preregistration and power analysis (https://doi.org/10.17605/OSF.IO/QV4YA).

## Results

### Confirmation of PFF-induced seeding of pSyn-positive aggregates

To confirm PFF-induced seeding, nigral slices were stained for TH and phosphorylated-synuclein (pSyn) from the remaining tissue of brains used for FSCV. We were unable to confirm seeding in animals used for uptake since the entire brain was used for that assay. We confirmed seeding in all animals used for FSCV and counted the number of pSyn-positive objects. At 4 months post-injection, as expected, we observed pSyn positive inclusions ipsilateral, but not contralateral, SN, in both dieldrin and vehicle/PFF groups (**Error! Reference source not found**.A-B). Consistent with previous results, dieldrin did not affect the number of pSyn-positive inclusions (**Error! Reference source not found**.C).^24^

### Developmental dieldrin exposure increases DA release in PFF-injected animals

FSCV was performed in striatal slices to determine if developmental dieldrin exposure affects evoked DA release or uptake in PFF-injected animals 4 months after PFF injection (Figure 4A-D). There was a significant increase in both peak DA concentration and upward velocity, a measure of DA release, in the dieldrin/PFF group compared to the vehicle/PFF group (*p*=0.0394 and *p*=0.0434, respectively) (Figure 4E,F). However, there was no significant difference in tau or downward velocity, which are measures of uptake (*p*=0.6435 and 0.5303 respectively) (Figure 4G,H). Calculated values are shown in Table 2. We verified that there was no difference in any metric on the contralateral side to confirm that dieldrin alone had no effect on DA release or uptake (Supplemental Figure 2A-D). We also compared ipsilateral to contralateral metrics in the vehicle/PFF group to and observed no significant effect of PFFs alone (Supplemental Figure 2E-H). Monomer injected animals in both the vehicle and dieldrin exposed groups showed similar outcomes on all FSCV metrics.

**Table 2.**
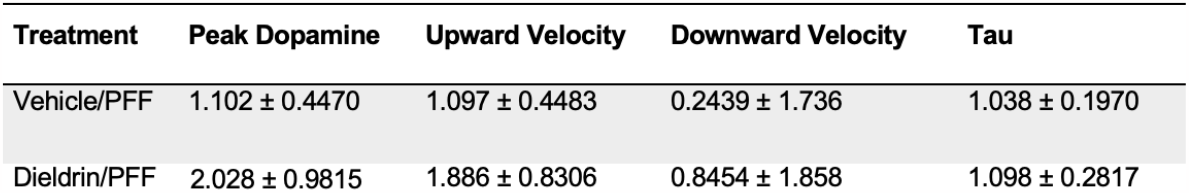
FSCV values (ipsilateral/contralateral) for vehicle/PFF and dieldrin/PFF groups (mean ± standard deviation)

### Developmental dieldrin exposure does not alter VMAT2 uptake velocity in PFF-injected animals

To determine if dieldrin exposure alters VMAT2 function in PFF-injected animals, uptake assays were performed at 4 months post-PFF injection. Somewhat surprisingly, there was no difference in VMAT2-mediated uptake velocity between the vehicle/PFF and the dieldrin/PFF groups ipsilateral to injection site (Figure 5). As expected, there was no difference in uptake contralateral to the injection site, showing that dieldrin alone had no effect on uptake velocity (Supplemental Figure 1A). In addition, uptake was equivalent between the ipsilateral and contralateral sides within the vehicle/PFF group, demonstrating no significant effect of PFFs alone (Supplemental Figure 1B). Observed uptake velocity was consistent with previously published values for VMAT2 uptake velocity in WT C57BL/6 mice. ^48^ Vehicle and dieldrin exposed animals injected with monomer showed similar VMAT2 uptake velocity in the hemisphere ipsilateral to the injection (vehicle/monomer: 7.020 ± 2.224 pmol/mg/min, n = 4; dieldrin/monomer: 5.460 ± 1.678 pmol/mg/min, n = 4).

## Discussion

### Developmental dieldrin exposure alters the dopaminergic response to synucleinopathy-triggered toxicity

Here, we demonstrate that developmental dieldrin exposure leads to an altered response to synucleinopathy and enhanced DA release in PFF-injected animals 4 months after PFF-injection (Figure 4). Specifically, we observed a dieldrin-related increase in DA release, but no change in DAT uptake by FSCV and no change in VMAT2-mediated uptake by uptake assay (Figure 5). These results are summarized in context with findings from our previous paper in Figure 6.^24^ These observed increases in DA release in dieldrin/PFF animals are consistent with our previously observed increase in DA turnover in PFF-injected animals developmentally exposed to dieldrin at 6 months post-PFF injection. If more DA is released, but DAT and VMAT2 uptake velocities are not changed, this could lead to increased cytosolic DA and the increased DA turnover that we previously observed at 6 month post-PFF injection.^39^ Importantly, our current FSCV and uptake data was collected at 4 months post PFF injection while our previous data showing increase striatal DA turnover was collected at 6 months post injection, suggesting that this enhanced DA release precedes enhanced toxicity.^24^ In addition, in our previous study, we reported the loss of striatal DA and its metabolites, DOPAC and HVA, at 2 and 6 months post-PFF injection that was not affected by dieldrin exposure.^24^ Together this suggests that synucleinopathy-induced DA loss occurs early and is not exacerbated by dieldrin, but the response to this DA deficit is affected by prior dieldrin resulting in enhanced DA release and an increase in DA turnover.

**Figure 3.**
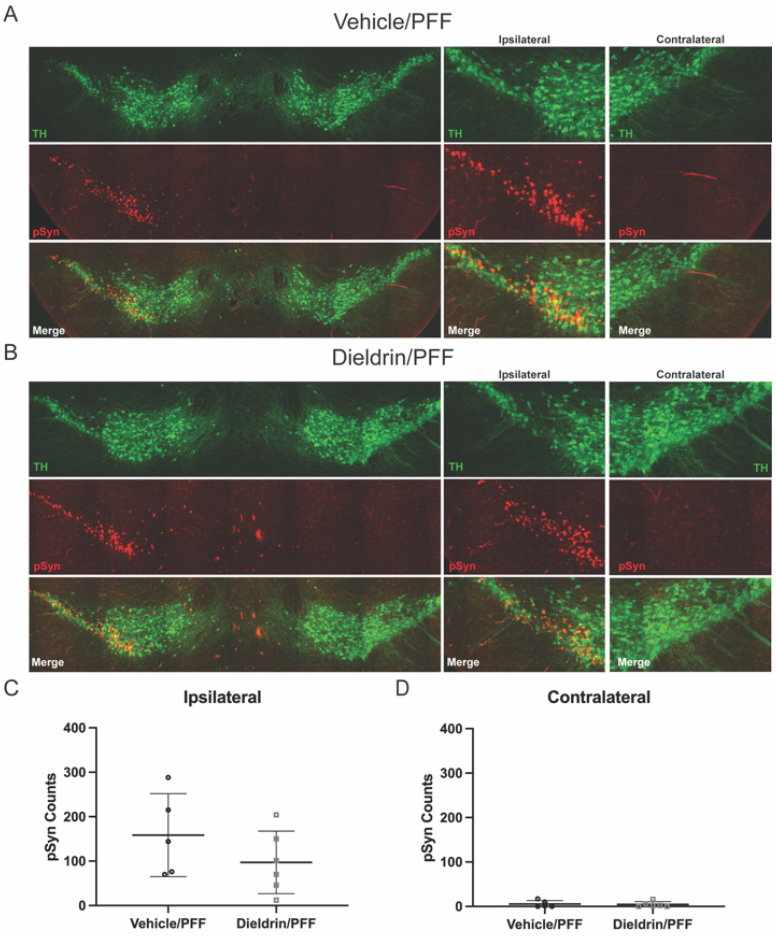
Confirmation of PFF-induced seeding in FSCV animals. A) Representative images from nigral tissue sections stained with TH (green) and pSyn (red) from a vehicle/PFF (A) and dieldrin/ PFF (B) animals 4 months post-PFF injection. C) pSyn counts in the SNpc show no effect of dieldrin on pSyn-positive objects in the SNpc ipsilateral to the PFF injection (*p=0*.*2441*). D) As expected, there were no pSyn-positive objects contralateral to the injection in either group of animals. All data are shown as mean +/- SD.

**Figure 4.**
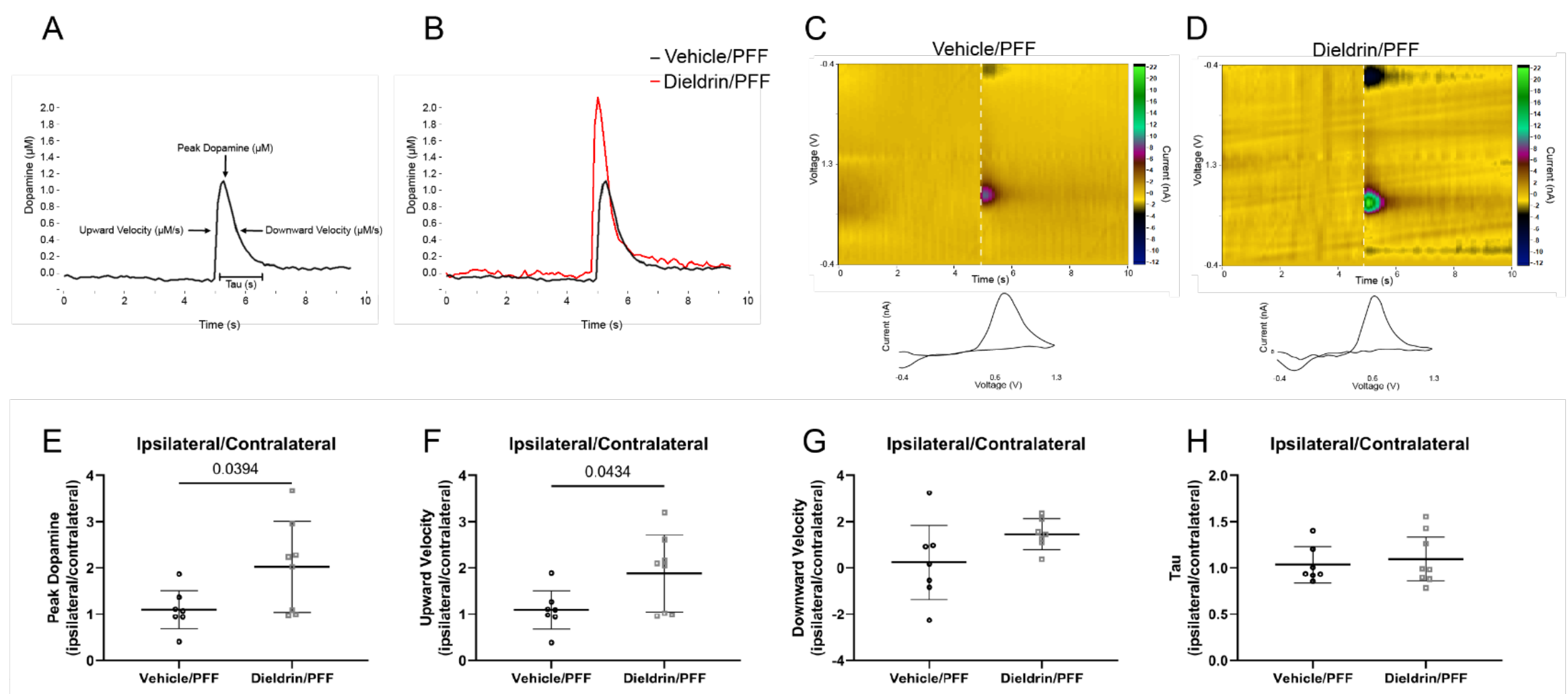
Dieldrin/PFFs increase peak dopamine and upward velocity in striatal tissues measured using FSCV. 4-months post-PFF injection, animals were sacrificed and FSCV was performed in dorsal striatum. A) Example dopamine versus time graph showing each quantified metric. B) Representative dopamine versus time graph for the groups vehicle/PFF (black) and dieldrin/PFF (red). C,D) Representative dopamine concentration vs time plot for (C) vehicle/PFF and (D) dieldrin/PFF following stimulation at t=5 secs. E-H) FSCV metrics represented as ipsilateral values normalized to contralateral values. E) Quantification of peak dopamine showed a significant dieldrin-related increase (*p*= 0.0394). F) Quantification of upward velocity showed a significant dieldrin-related increase (*p*= 0.0434). G) Quantification of downward velocity showed no significant effect of dieldrin (*p= 0*.*5303*). H) Quantification of tau showed no significant effect of dieldrin (*p= 0*.*6435*). Each individual data point represents a sum of 20 recordings per animal. All data shown as mean +/- SD.

**Figure 5.**
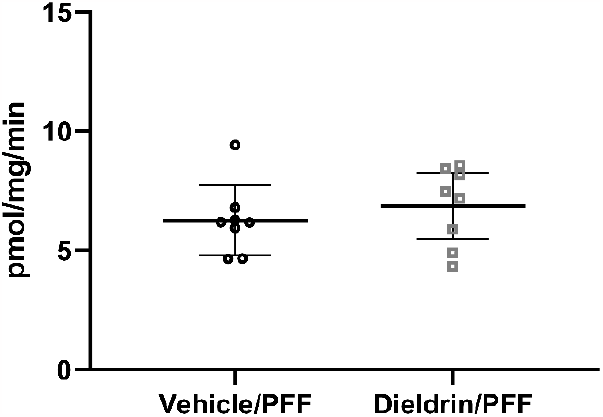
Dieldrin does not affect VMAT2 uptake velocity in PFF-injected male F1 offspring. There was no difference in uptake velocity ipsilateral to injection site 4 months post-PFF injection (*p*= 0.4524). All data shown as mean +/- SD.

**Figure 6:**
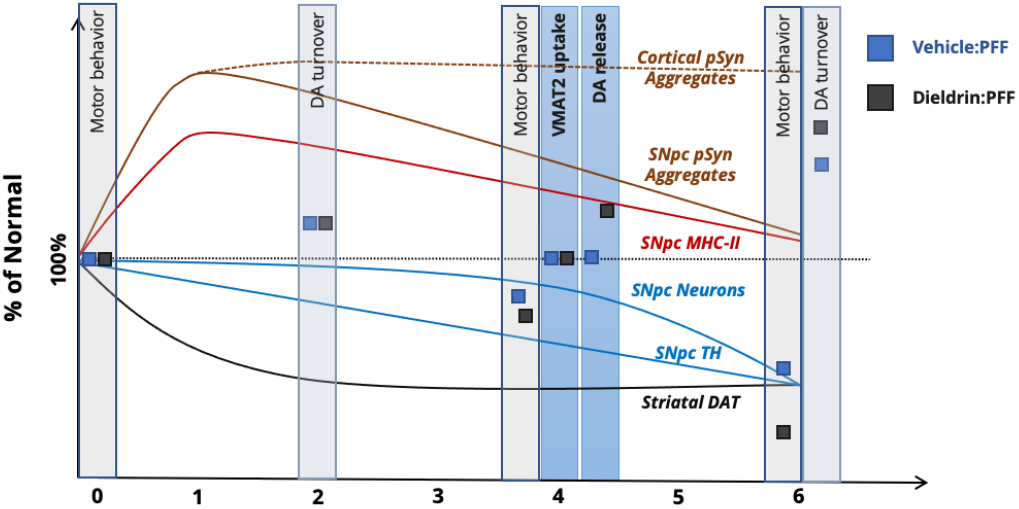
Summary of observed changes in the dieldrin PFF two hit model. Timelines show representative changes synuclein pathology, microglial activation, striatal loss, and nigral degeneration in the PFF model based on published literature, shown as the percent change in these markers compared to a saline/monomer injected mouse. Grey boxes indicate previous results from our lab in the dieldrin PFF two hit model. Blue boxes indicate FSCV and uptake results reported here at 4 months post-PFF injection. Blue and grey squares represent results from vehicle:PFF and dieldrin:PFF animals, respectively.

While FSCV has been utilized in α-syn knockout and α-syn overexpressing models, to our knowledge, this is the first study to perform FSCV in either the dieldrin or α-syn PFF model.^65–67^ These previous FSCV studies and other studies indicate a critical role for a-syn in DA release, synaptic vesicle fusion, vesicle trafficking, and regulation of synaptic vesicle pool size. Given that we did not observe any effect of dieldrin or PFF alone on DA release, dieldrin exposure appears to cause changes in the synaptic terminal that prime the nigrostriatal system for an exacerbated response to synucleinopathy (Supplementary Figure 2). This is consistent with our previous results demonstrating that this developmental exposure causes persistent epigenetic and transcriptomic changes in genes related to DAergic differentiation and maintenance in the midbrain and alters expression of neuroinflammatory genes in the striatum.^24,26^

Contrary to our hypothesis, we did not observe the PFF-induced decrease in DA release that we expected based on our and others previous data in PFF-injected mice showing loss of total striatal DA levels by 2 months post-PFF injection.^22,24^ This discrepancy is likely due to differences in methods and the underlying biology of DA neurons. HPLC measures total tissue DA levels from relatively large tissue punches, while FSCV measures extracellular DA only from the area immediately surrounding the electrode, so these measurements are not always aligned. Biologically, it is possible that while there is a significant PFF-induced loss of total tissue DA at 2 months post-PFF injection, reductions in DA release at these synapses is delayed relative to this loss even at 4 months post-PFF injection. This is consistent with a large body of data showing that most DA synapses within the striatum are silent and the majority of synaptic vesicles are located within the reserve pool rather than the readily releasable pool.^68–72^ Within the striatum of PFF-injected animals, we expect that only about a third to a half of terminals to be affected, so there is likely a pool of surviving neurons and vesicles within affected neurons to recruit to the readily releasable pool that allow the system to function. We previously observed only mild PFF-associated effects on motor behavior at this same 4 month post-PFF injection time point, suggesting that DA release is maintained even with about 45% loss of total striatal DA at 2 months post-PFF.^24^ In line with this, it is generally accepted that DA-related symptoms in human PD do not present until at least 30% of dopaminergic neurons in the nigrostriatal pathway are lost, suggesting that remaining neurons and striatal terminals are able to release sufficient quantities of DA despite a significant loss of total tissue DA levels.^73^

All together this suggests that the effects of dieldrin on PFF-induced deficits may be the result of an increased compensatory response to early DA loss in the striatum. Specifically, the observed increase in DA release may represent such a compensatory response to synucleinopathy that leads to greater toxicity in the long run due to resulting increases in cytosolic DA and resulting oxidative stress. ^74–83^ Early compensatory changes in the nigrostriatal system that precede degeneration are well-documented in human PD, multiple animal models of DA deficits, and more recently in a model of other monoaminergic (norephrinine) loss.^74–84.^ Thus, it is possible that developmental dieldrin exposure accelerates the toxic interplay between dysregulated α-syn and DA via early increases in compensatory mechanisms. Such a relationship between DA and α-syn is well-established and interfering with either can lead to a cycle of neurotoxicity where DA and α-syn interact and exacerbate the toxic effects of one another.^66,85–90^

### Developmental dieldrin exposure does not affect DAT- or VMAT2-mediated uptake in PFF-injected animals

The lack of significant effects on VMAT2 uptake velocity was surprising and inconsistent with our initial hypothesis. In the original developmental dieldrin study, it was demonstrated that developmental dieldrin exposure led to increases in striatal DAT and VMAT2 levels by western blot in both males and females and a male-specific increase in the DAT: VMAT2 ratio and a corresponding increase in DOPAC and DOPAC: DA at 12 weeks of age (the age at which we performed PFF injections).^25^ However, in our lab, we did not replicate these changes in DAT and VMAT2 levels at 12 weeks of age.^24^ Based on our previous western blot and HPLC results, we expected to see a relative increase in DAT function compared to VMAT2 function 4 months after PFF injection that was greater in animals developmentally exposed to dieldrin. A decrease in VMAT2 function relative to DAT function would lead to increased cytosolic DA and DA turnover and would explain our previously observed dieldrin-induced exacerbation of PFF-induced changes in DA turnover. ^24,25^

However, we did not observe any dieldrin related effect on VMAT2 uptake (Figure 5). It is possible that there is an effect on VMAT2 uptake velocity in the intact system that was not observed here due to experimental design and set up. Specifically, isolating synaptic vesicles for this assay removes them from their biological context and measures persistent changes due to overall changes in VMAT2 level or irreversible pharmacological effects.^44,48,91,92^ In addition, this assay involves homogenizing the entire hemisphere of the brain so effects specific to the striatum may be diluted by DAergic vesicles from areas of the brain not affected in our PFF model, including unaffected terminals within the striatum. Even within the striatum, we expect that only about a third to a half of terminals to be impacted. Thus, this assay may not have the sensitivity to detect a change in this small subpopulation of terminals. Unfortunately, technical limitations preclude us from performing this assay on unilateral striata from mouse brain.

It is also possible to increase the relative levels of DAT to VMAT2 function by affecting DAT function. Multiple prior studies show that both dieldrin and PFFs can impact DAT expression and function.^20,22,25,93^ Specifically, as discussed above, developmental dieldrin exposure was previously reported to lead to increases in striatal DAT levels at 12 weeks of age and changes in the DAT/VMAT2 ratio, but we did not observe the same effect in our previous study.^24,25^ A significant decrease in striatal DAT protein expression at 6 months post-PFF injection in C57BL/6 mice was observed, but not at 1- and 3-months post-injection.^22^ In Fischer 344 rats, DAT binding via PET imaging results shows a significant reduction as early as 2 months post-PFF injection with binding declining in a time-dependent manner.^93^ PFF-induced pathology is known to be slower and is less severe in mice compared to rats and may account partially account for this discrepancy in the relative timing of DAT decreases.^22,94^ Thus, it is possible that dieldrin-induced increases in striatal DAT function lead to a less severe loss of DAT following PFF-induced synucleinopathy and a relative increase of DAT to VMAT2 function. We observed an increase in total DA release, but no change in DA uptake in this study as measured by Tau and downward velocity from FSCV data (Figure 4). Thus, it is possible that if more DA is released, but neither DAT- or VMAT2-mediated uptake velocity is the same, the observed increase in turnover is due to extracellular metabolism of DA rather than intracellular breakdown.

Despite these caveats regarding VMAT2 and DAT uptake, this new data is consistent with our previous results in this two-hit model showing no increase in α-syn pathology but an enhanced response to synucleinopathy.^24^ Our results are also consistent with the idea of silent neurotoxicity, where the effects of early life exposures are unmasked by challenges later in life, the cumulative effects of exposures over the lifespan, or the effects of aging.^95,96^ In such a paradigm, developmental exposure to dieldrin primes the nigrostriatal striatal system in male offspring to have an exacerbated response to synucleinopathy induced by α-syn PFFs in the absence of observable changes in typical markers of nigrostriatal dysfunction and degeneration. In support of this, our previous studies identified persistent epigenetic and transcriptomic changes in genes related to DAergic differentiation and maintenance in the midbrain and altered expression of neuroinflammatory genes in the striatum at 12 weeks of age following developmental dieldrin exposure.^24,26^ In a parallel study, we are also tracking the longitudinal patterns of dieldrin-induced epigenetic changes across the timeline of this entire two-hit model from birth to 9 months of age to determine if dieldrin alters the trajectory of epigenetic changes across the lifespan.

### A model of dieldrin-induced increases in susceptibility to synucleinopathy

Based on the results reported here and our previous studies, we propose a model for how developmental dieldrin exposure leads to increased susceptibility to synucleinopathy-induced deficits in motor behavior.^24–26^. In this model, exposure to dieldrin occurs during prenatal and postnatal development. The half-life of dieldrin in mouse brain is less than a week, so no detectable dieldrin remains in the brain of F1 offspring by a few weeks after weaning. ^20,25,97^ When dieldrin is present in the developing brain, it is thought to act on developing DA neurons by inhibiting GABAA receptor-mediated chloride flux, resulting in increased neuronal activity.^98–103^ Based on previous results, we propose that this net increase in neuronal activity modifies the dopamine system through persistent changes in epigenetic mechanisms, leading to dysregulation of genes important for dopamine neuron development and maintenance, as well as changes in the neuroinflammatory system in the striatum.^24,26^ These changes alter the response of the nigrostriatal system to future insults via alterations in striatal dopamine synapses that manifest after induction of synucleinopathy by PFF injection as an increase in compensatory mechanisms triggered by striatal DA loss that precedes an increase in DA turnover. ^24^(Figure 4). Taken together, these results suggest that exploring dieldrin-induced changes that produce this high susceptibility state are critical to advancing our understanding of how exposures contribute to increased risk of PD and underscore the need to study PD-related exposures across the lifespan, particularly during sensitive periods of neurodevelopment.

## Data Availability

This study was preregistered with Open Science Framework: https://doi.org/10.17605/osf.io/qv4ya All data acquired and analyzed for this study are available the Figshare data repository: <u>10.6084/m9.figshare.23546586</u>

## CRediT Author Statement

**Sierra L. Boyd:** Investigation, Software, Formal Analysis, Writing – Original Draft, Writing – Review & Editing, Visualization, Project Administration; **Nathan C. Kuhn:** Methodology, Investigation, Project Administration; **Joseph R. Patterson:** Investigation, Validation, Writing – Review & Editing; **Anna C. Stoll:** Investigation; **Sydney A. Zimmerman:** Investigation; **Mason R. Kolanowski:** Investigation; **Joseph J. Neubecker:** Investigation; **Kelvin C. Luk:** Resources; **Eric S. Ramsson:** Conceptualization, Investigation, Supervision, Writing – Review & Editing; **Caryl E. Sortwell:** Conceptualization, Investigation, Supervision, Writing – Review & Editing; **Alison I. Bernstein:** Conceptualization, Supervision, Data Curation, Writing – Review & Editing, Funding Acquisition

## Supplementary figures

**Supplementary Figure 1.**
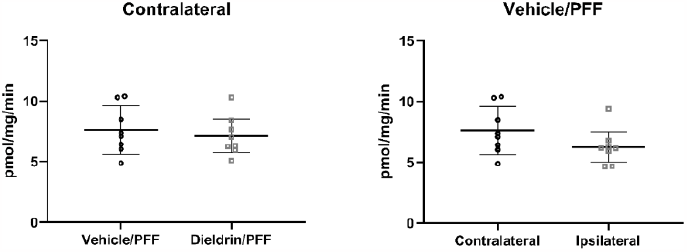
Developmental dieldrin exposure or PFF injection alone do not affect VMAT2-mediated vesicular uptake. A) There was no effect of dieldrin on VMAT2-mediated uptake on the non-injected contralateral side (*p=0*.*5972*). B) There was no effect of PFF on uptake from comparing ipsilateral to contralateral in vehicle/PFF animals (*p=0*.*0871*). Each data point represents an individual animal. All data shown as mean +/- SD.

**Supplementary Figure 2.**
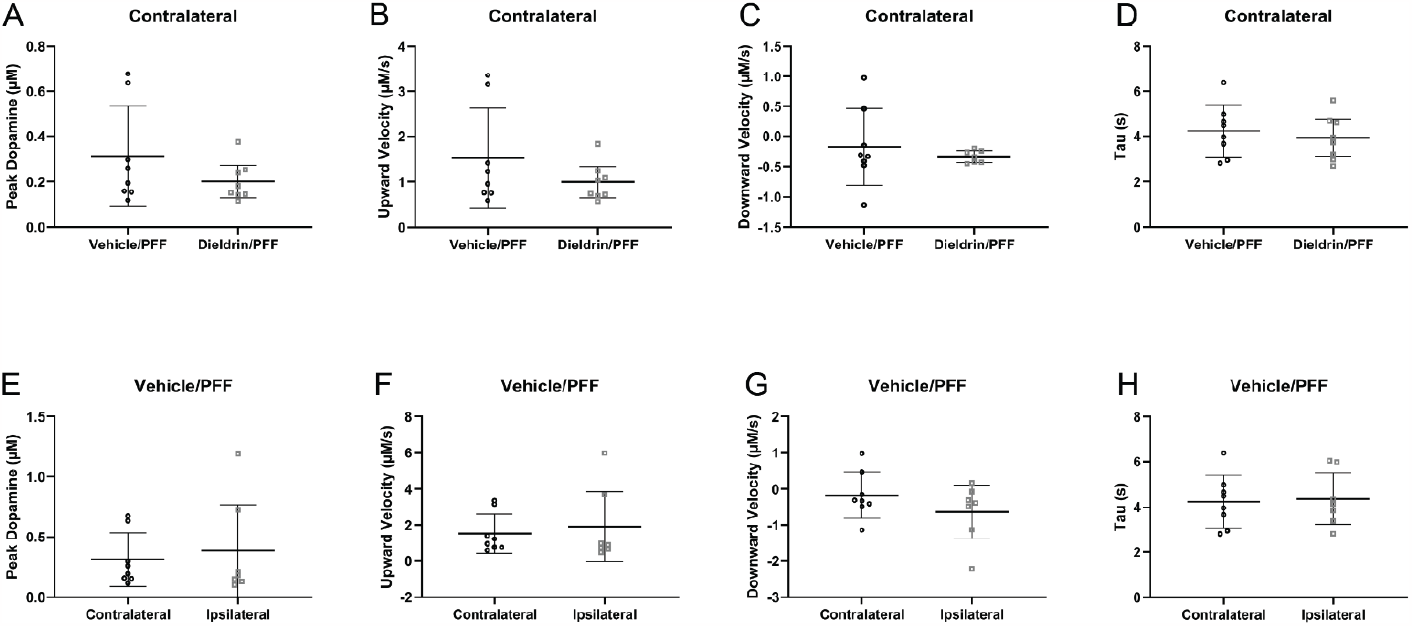
Developmental dieldrin exposure or PFF injection alone do not affect FSCV metrics. A-D) There was no dieldrin-related effect on the contralateral side on peak dopamine, upward velocity, downward velocity, or tau between vehicle/PFF versus dieldrin/PFF, indicating that dieldrin alone has no effect on FSCV metrics at this time point (*p=0*.*2033, 0*.*2189, 0*.*5133*, and *0*.*5736*, respectively). E-H) There was no PFF-related effect on peak dopamine, upward velocity, downward velocity, or tau between the contralateral versus ipsilateral striata, indicating that PFF injection alone has no effect on FSCV metrics at this point (*p=0*.*6708, 0*.*6562, 0*.*2395*, and *0*.*8440* respectively). Each individual data point represents a sum of 20 recordings per animal. All data shown as mean +/- SD.

**Supplementary Table 1.**
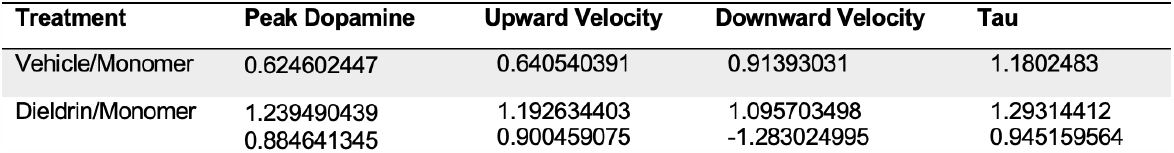
FSCV values (ipsilateral/contralateral) for each individual animal in vehicle/monomer and dieldrin/monomer groups.

